# Stress-Dependent Conformational Changes of Artemin: Effects of Heat and Oxidant

**DOI:** 10.1101/2020.02.12.946236

**Authors:** Z. Takalloo, Z. Afshar Ardakani, B. Maroufi, S. Shirin Shahangian, R. H. Sajedi

## Abstract

Artemin is an abundant thermostable protein in *Artemia* embryos and considered as a highly efficient molecular chaperone against extreme environmental stress conditions. The dynamic conformational properties of artemin appear to play a critical role in its biological activities. In this study, we have investigated the conformational transitions and functional changes of artemin under heat and oxidative stress to find some evidence of the relationship between the structure and function of artemin. The tertiary and quaternary structures of artemin have been evaluated by fluorescence measurements, protein cross-linking analysis, and dynamic light scattering. Based on the structural analysis, artemin showed irreversible substantial conformational lability in response to heat and oxidant which mainly mediated through the hydrophobic interactions and dimerization of the chaperone. In addition, the chaperone-like activity of the heated and oxidized artemin was examined using lysozyme refolding assay and the experiments showed that although both factors, i.e. heat and oxidant, at specific levels improved artemin potency, simultaneous incubation with both stressors significantly triggered the activation of artemin. Moreover, the heat-induced dimerization of artemin was found to be the most critical factor for its activation. It was suggested that oxidation presumably acts through stabilizing the dimer structures of artemin through formation of disulfide bridges between the subunits and strengthens its chaperoning efficacy. Accordingly, it was proposed that artemin probably exists in a monomer–oligomer equilibrium in *Artemia* cysts and environmental stresses and intracellular portion of protein substrates may shift the equilibrium towards the active dimer forms of the chaperone.

**STATEMENT OF SIGNIFICANCE:** There are a number of reports in which the chaperone-like activity of artemin, as a stress protein, has been confirmed *in vivo* and *in vitro.* Nonetheless, the details of structural changes of the protein have not been fully discovered yet. In the present work, we focused on conformational properties of artemin from *A. urmiana* upon exposing to heat and oxidation, by using various structural and functional analysis in order to predict the mechanisms of artemin’s activation. Notably, this is the first document on reporting the structural transitions of the chaperone in stress conditions.

## INTRODUCTION

Artemin is a stress protein of encysted *Artemia* embryos, representing about 10% of the soluble cellular proteins, but it is almost completely absent from nauplius larvae (1). Due to high structural stability, abundance and chaperone function, artemin probably contributes to cyst stress resistance (2). Artemin monomers consist of 229 amino acid residues and exhibit a molecular mass of 26 kDa and assemble into rosette-like oligomers of ∼600–700 kDa consist of 24 monomer subunits (2, 3). Artemin and ferritin, an iron storage protein, show a degree of homology in both sequence and structure, although artemin contains 45–50 extra residues compared to ferritin (1,3–5). In contrast, artemin fails to bind iron due to the extension of carboxyl terminal of the protein monomers which suggests to have a role in its chaperone function (6, 3). It is very heat stable and associates with RNA at elevated temperatures, suggesting protection of RNA (7).

Previous researches revealed a chaperone-like activity for artemin which is responsible for protecting transfected cells and preventing protein aggregation *in vitro* (8). Our previous computational and experimental studies on artemin from *Artemia urmiana* (Urmia Lake, Iran) also confirmed the chaperone potency of artemin in suppressing protein aggregation (3,6,9). Besides, artemin was capable to protect proteins and cells against different stress conditions such as oxidant, cold and salt (10, 11). These studies proposed that artemin probably binds to protein substrates *via* hydrophobic interactions. Artemin is a histidine/cysteine-rich protein and due to the high content of cysteines and their distributions, it was suggested that the oxidative state of cysteines moderates the redox-regulated activity of artemin (1, 12). Therefore, in our recent study, the modification of cysteine residues revealed that the chaperone potency of artemin is greatly depended on the formation of intermolecular disulfide bonds introducing artemin as a redox-dependent chaperone (13).

Our suggestion is that conformational changes of artemin are responsible for its chaperone function upon exposing to stress conditions. In fact, these structural modifications may change the surface hydrophobicity of the chaperone, which result in hydrophobic interactions with target proteins. Accordingly, in the present study, The tertiary and quaternary structures of artemin under heat and oxidative stresses have been evaluated by intrinsic and ANS fluorescence measurements, protein cross-linking analysis, and dynamic light scattering (DLS). In addition, the chaperone function of artemin was investigated under both stress conditions using dilution-induced aggregation assay of lysozyme.

## MATERIALS AND METHODS

### Chemicals

Glycine, Tris, NaCl, sodium dodecyl sulfate (SDS), imidazole, kanamycin and Isopropyl-β-D-thiogalactopyranoside (IPTG) were purchased from Bio Basic Inc. (Markham, Ontario, Canada). Peptone and yeast extract were provided by Micromedia Trading House Ltd (Pest, Hungary). AgNO_3_ and 8-anilino-l-naph-thalene sulfonic acid (ANS) were purchased from Sigma–Aldrich (St. Louis, MO, USA). Nickel-nitrilotriacetic acid agarose (Ni-NTA agarose) was provided by Qiagen (Hilden, Germany). Chicken egg white lysozyme, dithiothreitol (DTT), glutaraldehyde, H_2_O_2_, glycerol, and all other chemicals were obtained from Merck (Darmstadt, Germany). Protein standard marker was purchased from Thermo Fisher Scientific (Waltham, MA, USA).

### Expression and purification of recombinant artemin

pET28a encoding artemin from *Artemia urmiana* was provided (3) and protein expression was carried out in *Escherichia coli* BL21 (DE3) cells as previously described (9). Purification of the His-tagged protein was carried out using Ni-NTA agarose column. Bound proteins were eluted with a buffer containing 50 mM NaH_2_PO_4_, 300 mM NaCl, and 250 mM imidazole, pH 8.0. Aliquots of the eluted protein were taken, followed by dialysis against the phosphate buffer overnight at 4°C. The protein concentrations were determined using Bradford’s method and BSA as standard (14).

### Protein treatments

#### Heated artemin (H-artemin)

Purified artemin was incubated at temperature range 25-80°C for 20 min followed by incubation at room temperature for 20 min before measurements.

#### Oxidized artemin (O-artemin)

Purified artemin was incubated with 0-160 mM H_2_O_2_ for 6 h at 0°C in dark condition before measurements.

#### Heated oxidized artemin (HO-artemin)

Purified artemin was incubated with 0-100 mM H_2_O_2_ for 6 h at 0°C in dark condition, then incubated at temperature range 25-70°C for 20 min followed by incubation at room temperature for 20 min before measurements.

### Conformational analysis

#### Intrinsic fluorescence measurements

Intrinsic aromatic fluorescence measurements were performed using a LS-55 fluorescence spectrometer (Perkin-Elmer, USA). Samples containing H-artemin, O-artemin, and HO-artemin in 50 mM phosphate buffer, pH 7.2 were used. The excitation wavelength was set at 280 nm, and the emission spectra were monitored in the wavelength range of 300–400 nm. Excitation and emission bandwidths were 10 nm. Protein concentration for intrinsic fluorescence measurements was about 0.1 mg/mL.

#### ANS fluorescence measurements

Samples containing H-artemin, O-artemin, and HO-artemin in 50 mM phosphate buffer, pH 7.2 were used. The excitation wavelength was set at 380 nm and emission spectra were taken from 400 to 600 nm by using the fluorescence spectrometer. Excitation and emission slits were set at 10 nm. Protein concentration for ANS measurements was about 0.25 mg/mL and ANS was added at a final concentration of 30 µM.

To investigate whether stress-induced conformational changes of artemin was reversible, the heated and oxidized artemin were kept at 4°C for 3 h and 48 h, then their intrinsic and extrinsic fluorescence were measured. In the case of oxidized proteins, for elimination of hydrogen peroxide, catalase (0.5 µg/mL) was added in the protein solution, followed by incubating the samples at 4°C.

### Protein cross-link analysis by SDS-PAGE

H-artemin, O-artemin, and HO-artemin at final concentration of about 0.2 mg/mL were used. Glutaraldehyde (0.5%) was added to the all protein treatments and the reactions were terminated after 1 min by addition of 200 mM Tris-HCl, pH 8.0 at final concentration. In the case of oxidized samples, catalase (0.5 µg/mL) was added to decompose the hydrogen peroxide in the solution before addition of glutaraldehyde. The protein cross-linking were analyzed using 10% non-reducing SDS-PAGE. The incubated artemin at 25°C was used as control.

In addition, to check whether the dimerization of artemin, especially in the case of heated artemin, is dependent on disulfide bridges, DTT (30 mM at final concentration) was added to artemin (0.1 mg/mL), then the samples were incubated at 25 and 60 °C for 20 min followed by incubation at room temperature for 20 min. Dimerization of the protein was checked using 10% non-reducing SDS-PAGE with and without DTT and glutaraldehyde as the reducing and cross-linking agent, respectively.

### DLS analysis

The size analysis of the H-artemin (0.05 mg/mL) at 25-80°C was carried out using dynamic light scattering by Zetasizer Nano ZS instrument (Malvern Instruments Ltd., Malvern, Worcestershire, UK).

### Chaperone-like activity assay

#### Preparation of denatured reduced lysozyme

Denatured reduced lysozyme (10 mg/mL) was prepared by diluting appropriate volume of the enzyme in a denaturation buffer containing 6 M GdmCl and 40 mM DTT in 50 mM potassium phosphate buffer, pH 7.1. The sample was incubated at room temperature for a period of 12 hours to yield a fully reduced denatured enzyme. The denatured reduced lysozyme was used for refolding studies.

#### Aggregation accompanying refolding

Refolding of lysozyme was initiated after 50-fold rapid dilution of the denatured enzyme solution (10 mg*/*mL) with a refolding buffer (5 mM GSH, 5 mM GSSG in 50 mM potassium phosphate buffer, pH 8.5) containing artemin at final concentrations of 0.5 and 1 µg/mL. The final concentrations of the redox reagents were in the range of the optimum concentrations suggested by a previous report (15). The final concentration of lysozyme was 0.2 mg/mL in the buffer. The kinetic of refolding was recorded by monitoring light scattering at 400 nm at room temperature for 2 min using a microplate spectrophotometer (µQuant, BioTek, USA). As a control, the denatured lysozyme was refolded as described without artemin. In these experiments, H-artemin (25-80°C), O-artemin (0-100 mM H_2_O_2_), and HO-artemin (25-100 mM H_2_O_2_, 25-70°C) were used.

## RESULTS

### Temperature-dependent structural changes in artemin

Intrinsic fluorescence was monitored at 330 nm with 280 nm excitation to probe changes in tertiary structures of H-artemin (Figure 1A). The results indicated that when the temperature increased up to 30°C, the fluorescence intensity decreased sharply, and the lowest emission intensity was recorded for the heated protein at 80°C (Figure 1A). Besides, the wavelength of maximum emission (λ_max_) did not show any shift.

**Figure 1.**
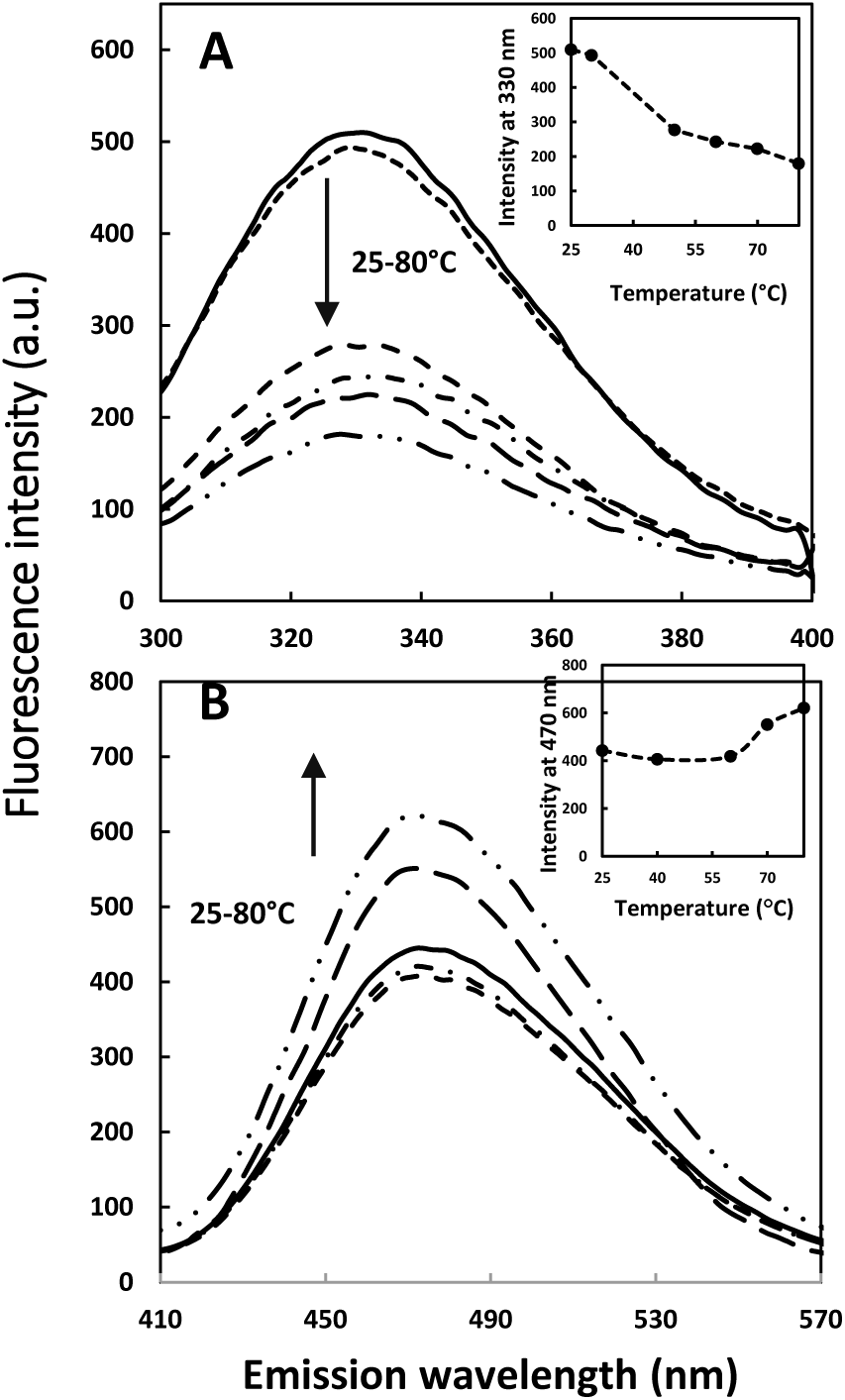
Intrinsic (A) and extrinsic (B) fluorescence emission intensity of artemin at different temperatures. A) Trp fluorescence emission of artemin was measured by incubating artemin (0.1 mg/mL in 50 mM phosphate buffer, pH 7.2) at 25-80°C for 20 min, followed by cooling down at room temperature for 20 min. The fluorescence intensity was recorded with the excitation wavelength at 280 nm. The inset is temperature dependence of the fluorescence intensity of artemin at 330 nm. B) Fluorescence emission intensity of ANS in the presence of heated artemin. Samples of artemin (0.25 mg/mL in 50 mM phosphate buffer, pH 7.2) were incubated at 25-80°C for 20 min, and cooled down at room temperature, then ANS (30 µM at final concentration) was added and fluorescence intensity was measured with the excitation wavelength at 380 nm. The insets indicate temperature dependence of the fluorescence intensity of artemin at 470 nm.

The fluorescence of 30 µM ANS in the presence of H-artemin was also monitored at 470 nm with 380 nm excitation (Figure 1B). ANS binding showed that the fluorescence intensity did not change considerably from 25 to 60°С. In contrast, a sharp emission peak was monitored at 470 nm for the heated proteins incubated at temperatures 70-80°C.

Glutaraldehyde cross-linking of artemin followed by SDS-PAGE revealed that upon increasing temperatures from 25 to 50°C, the band corresponding to dimeric form of artemin around 54 kDa slightly strengthen and after that, it was reduced at 60 °C, and finally completely disappeared at 70 and 80 °C (Figure 2A). Besides, the dimeric band disappeared following incubation of artemin with DTT before heat treatments under both cross-linking and non-cross-linking conditions showing that induced dimerization of H-artemin is dependent on the formation of disulfide bond(s) (Figure 2B).

**Figure 2.**
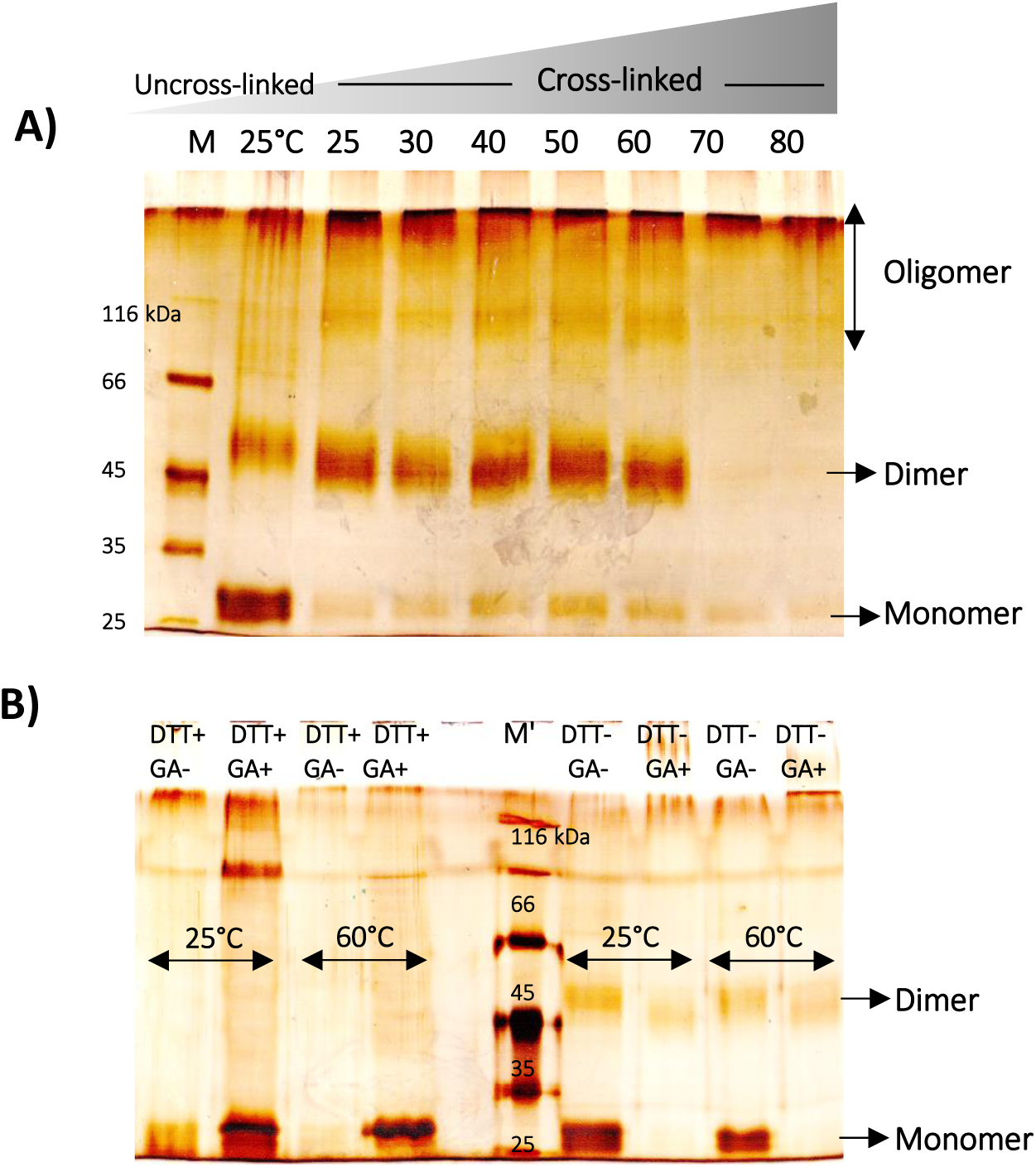
Protein cross-link analysis of heated artemin by non-reducing SDS-PAGE. A) Glutaraldehyde cross-linking of H-artemin. Artemin (0.2 mg/mL) was incubated at 25-80°C for 20 min, then cooled down at room temperature for 20 min. 0.5% glutaraldehyde was added and the reactions were terminated after 1 min by addition of 200 mM Tris-HCl, pH 8.0. The uncrossed artemin incubated at 25°C was used as control. B) The heated protein fractions treated with(out) DTT as the reducing agent. 30 mM DTT was added to artemin (0.1 mg/mL), followed by incubation of the samples at 25 and 60°C for 20 min. The (un)cross-linked (GA^(-)+^) (non)reduced (DTT^(-)+^) artemin was analyzed using 10% non-reducing SDS-PAGE. M; protein standard marker.

DLS measurement confirmed that the average size diameter of the heated chaperone increased upon enhancement of temperature (Figure 3) and accordingly, the largest size was detected at 70-80°C. The larger size distribution of H-artemin can be explained with its ability to form dimeric and oligomeric species as also indicated by SDS-PAGE (Figure 2A). All results showed that the conformational changes in artemin occur upon increasing temperature.

**Figure 3.**
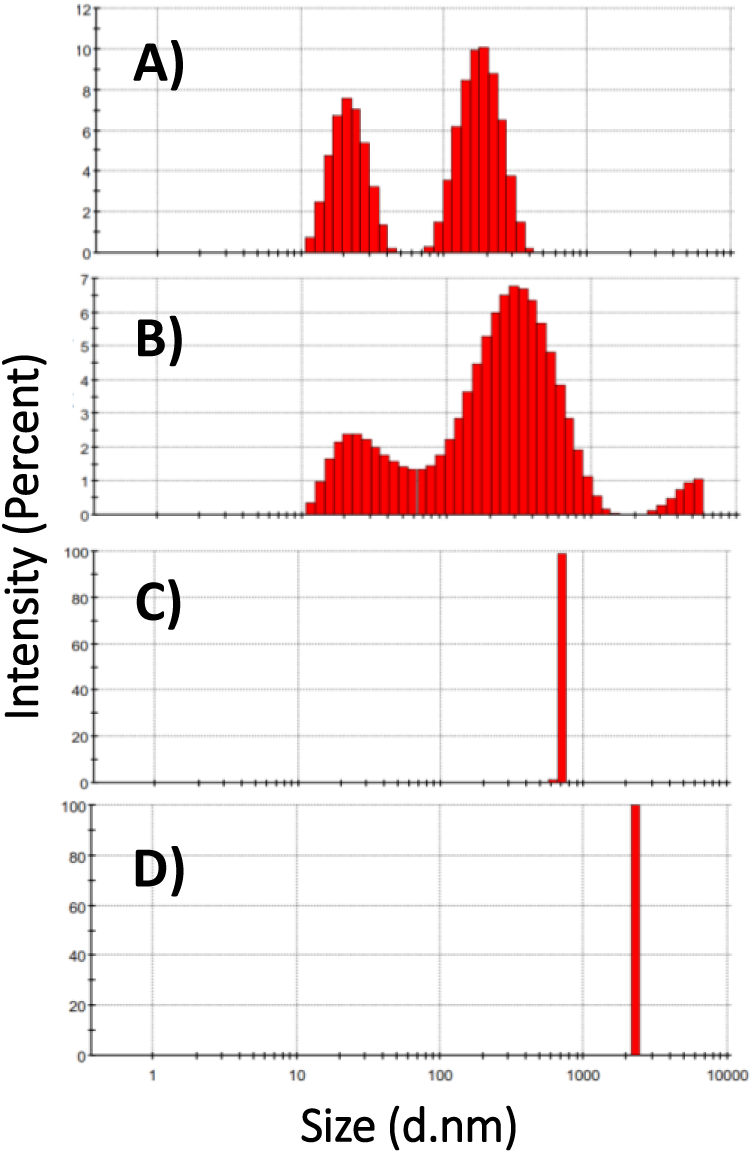
DLS analysis of heat-treated artemin. Artemin was incubated at 25 (A), 50 (B), 70 (C), 80 (D) °C for 20 min, then kept at room temperature for 20 min and size distribution analysis of H-artemin was performed.

### Oxidative-dependent structural changes in artemin

As depicted in Figure 4A, fluorescence emission maximum (λ_max_), as well as the fluorescence intensity of protein samples, was influenced by increasing the oxidant concentration. The slight decrease in fluorescence at 332 nm was monitored with increasing the concentration of oxidant from 2.5 to 40 mM, followed by a considerable decline in fluorescence intensity for O-artemin with 80-160 mM H_2_O_2_. Besides, λ_max_ shifted to longer wavelengths, from 330 to 337 nm (Figure 4A).

**Figure 4.**
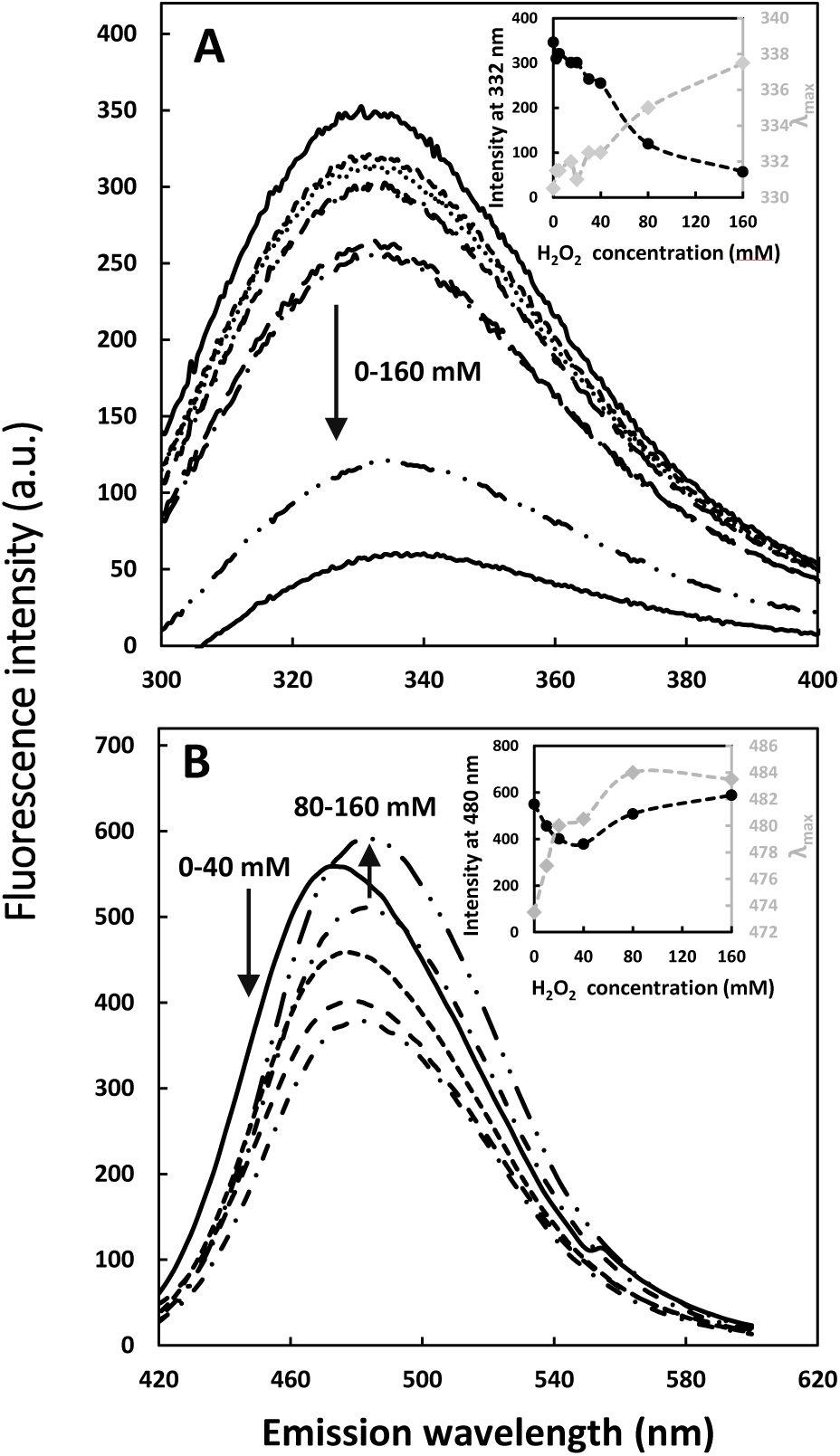
Intrinsic (A) and extrinsic (B) fluorescence emission intensity of artemin at different H_2_O_2_ concentrations. A) Intrinsic fluorescence emission of artemin was measured by incubating artemin (0.1 mg/mL in 50 mM phosphate buffer, pH 7.2) at 0-160 mM H_2_O_2_ for 6 h at 0°C in dark, then the fluorescence intensity was recorded with the excitation wavelength at 280 nm. The inset is the oxidant dependence of the fluorescence intensity of artemin at 330 nm. B) ANS fluorescence emission spectra was monitored after addition of 30 µM ANS to O-artemin (0.25 mg/mL in 50 phosphate buffer, pH 7.2). The excitation wavelength was 380 nm. The inset shows the oxidant dependence of the fluorescence intensity of artemin at 480 nm and also shifting the maximum emission wavelength (λ_max_) of the oxidized chaperone from 474 to 484 nm.

Extrinsic fluorescence using ANS showed a two-state process. The O-artemin with lower concentrations of H_2_O_2_ (0-40 mM), showed slight intensity decreases, but the intensity was enhanced at higher concentrations of oxidant (80-160 mM). As depicted in Figure 4B, a 10 nm red shift of the emission peak position of O-artemin is seen when spectra at lower oxidant contents (0-40 mM, ∼ 474 nm) are compared to those at higher oxidant concentrations (80-160 mM, ∼ 484 nm).

SDS-PAGE analysis of cross-linked artemin showed that the band corresponding to dimeric form of O-artemin gradually strengthened upon increasing the concentration of H_2_O_2_ from 0 to 100 mM, and it was weakened at 160 mM H_2_O_2_ (Figure 5).

**Figure 5.**
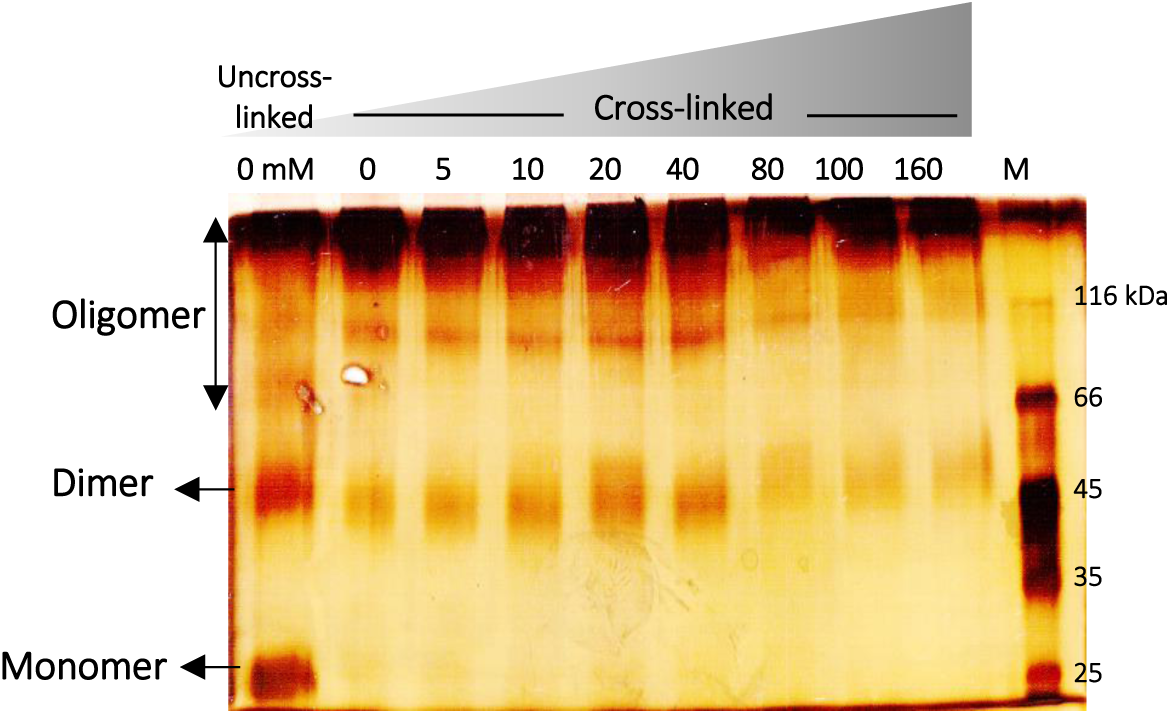
Protein cross-link analysis of oxidized artemin with various H_2_O_2_ concentrations by SDS-PAGE. Purified artemin (0.2 mg/mL) was incubated with 0-160 mM H_2_O_2_ for 6 h at 0°C in dark condition. Before the samples were loaded on 10% non-reducing SDS-PAGE, a trace amount of catalase (0.5 µg/mL) was added to the protein solutions to decompose the remaining H_2_O_2_. M; protein standard marker.

### Suggested place for Figure 5 Structural changes of artemin under heat and oxidative conditions

To evaluate the influence of the two stressors, i.e. heat and oxidant, on protein structure, artemin was treated with 0-100 mM H_2_O_2_, followed by exposure to 50 and 70°C. Intrinsic fluorescence measurements showed that under both temperature incubations, the intensity declined gradually by increasing the oxidant concentrations from 0 to 100 mM (Figure 6A). Besides, ANS fluorescence indicated that the fluorescence intensity did not change considerably for HO-artemin incubated with 10-100 mM H_2_O_2_ at elevated temperatures (Figure 6B). This trend was not similar to those observed for the individual oxidant treatments (Figure 4B).

**Figure 6.**
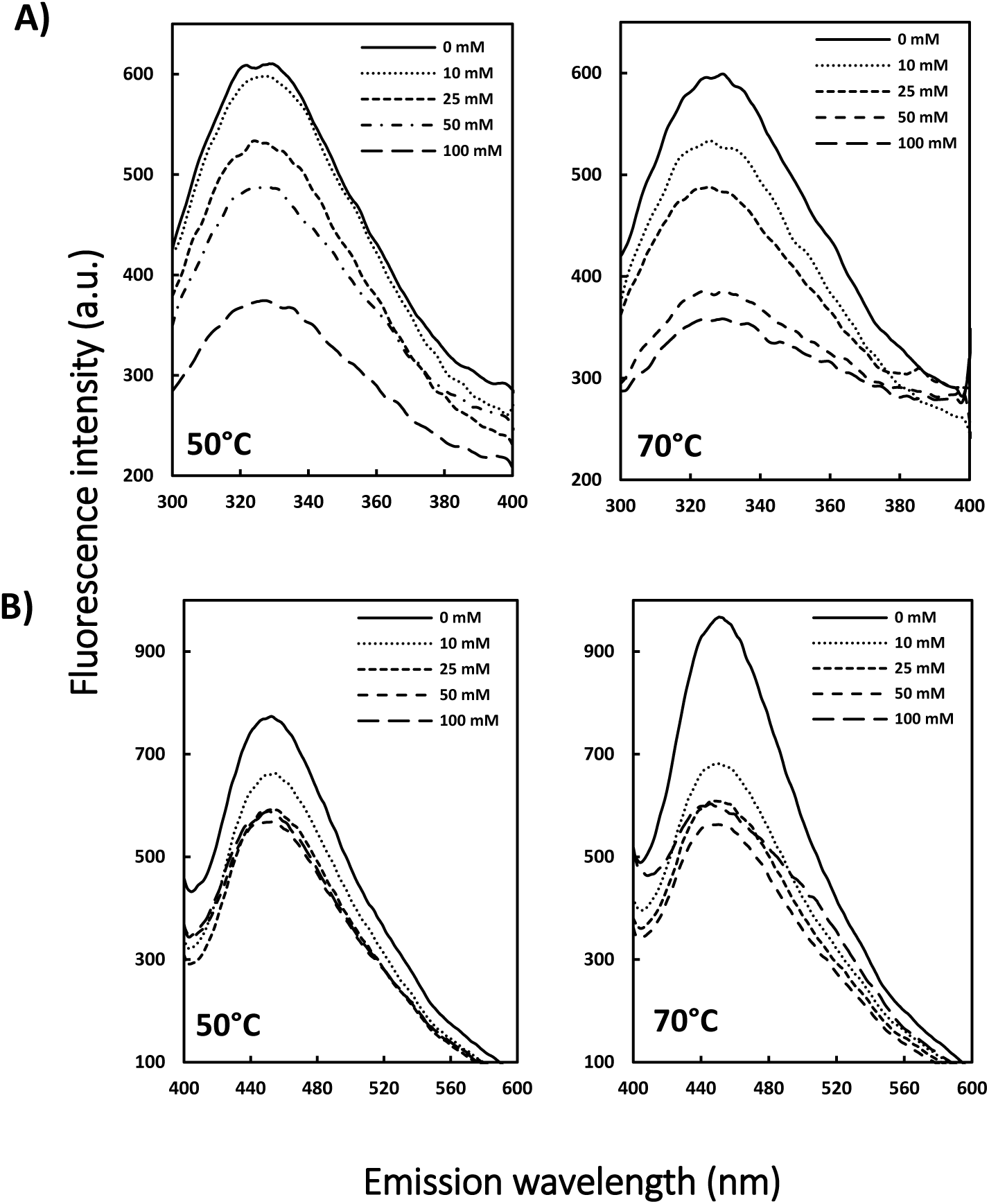
Intrinsic (A) and extrinsic (B) fluorescence emission intensity of heated oxidized artemin. A) Intrinsic fluorescence emission of artemin was recorded by incubating artemin (0.1 mg/mL in 50 mM phosphate buffer, pH 7.2) with 0-100 mM H_2_O_2_ for 6 h at 0°C in dark, followed by incubating the samples at 50 and 70°C for 20 min, then cooling down at room temperature. The fluorescence intensity was recorded with the excitation wavelength at 280 nm. B) Fluorescence emission intensity of ANS (30 µM at final concentration) was measured in the presence of HO-artemin (0.25 mg/mL) with the excitation wavelength at 380 nm.

In a second approach, the change in oligomerization state of HO-artemin was examined by SDS-PAGE (Figure 7). The bands representing the dimeric forms of HO-artemin were clearly strengthened along with enhancement of the oxidant concentrations.

**Figure 7.**
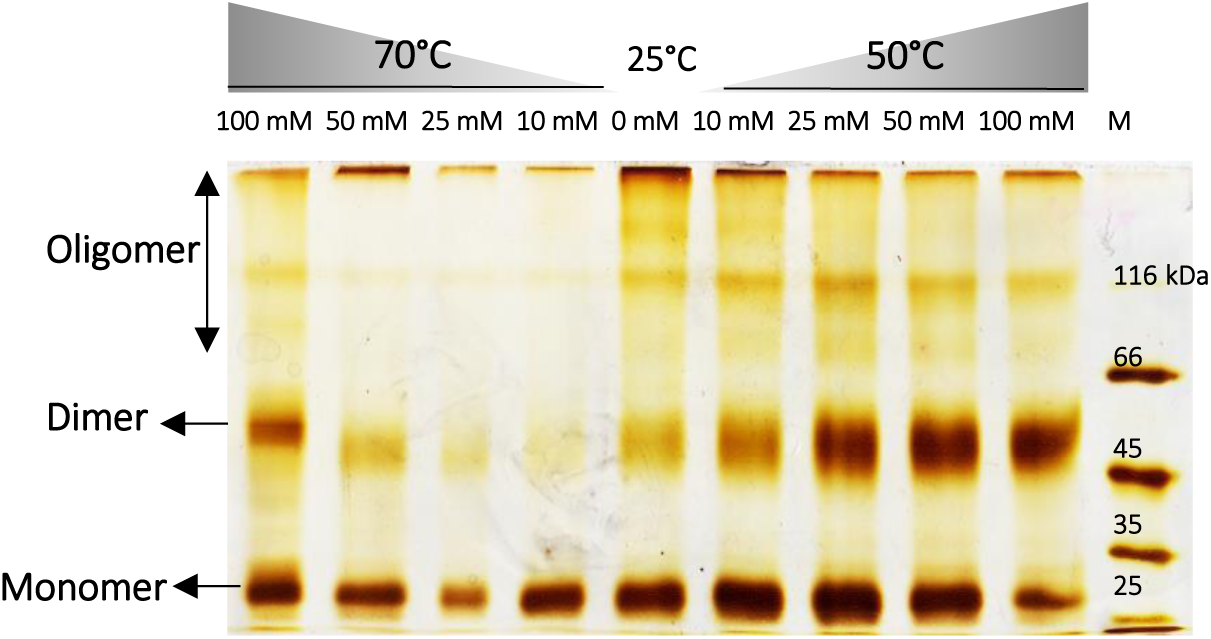
Cross-link analysis of heated oxidized artemin by SDS-PAGE. Purified artemin (0.2 mg/mL) was incubated with 0-100 mM H_2_O_2_ for 6 h at 0°C in dark condition. Then the samples were subjected to temperatures of 50 and 70°C for 20 min and cooled down at room temperature. Before the samples were loaded, catalase (0.5 µg/mL) was added to the solutions to decompose the remaining H_2_O_2_.

### Structural changes of artemin under stress conditions are irreversible

To determine whether the temperature/oxidation-dependent transition of artemin is reversible, artemin was incubated at elevated temperatures (H-artemin), high oxidant concentrations (O-artemin), and both heat and oxidant conditions (HO-artemin), followed elimination of the heat and oxidant and keeping the samples at 4°C for 48 h before fluorescence measurements (Figure 8). The results showed that fluorescence intensity did not change after 48 h. In fact, the structure of artemin modified irreversibly upon exposing to the stressors.

**Figure 8.**
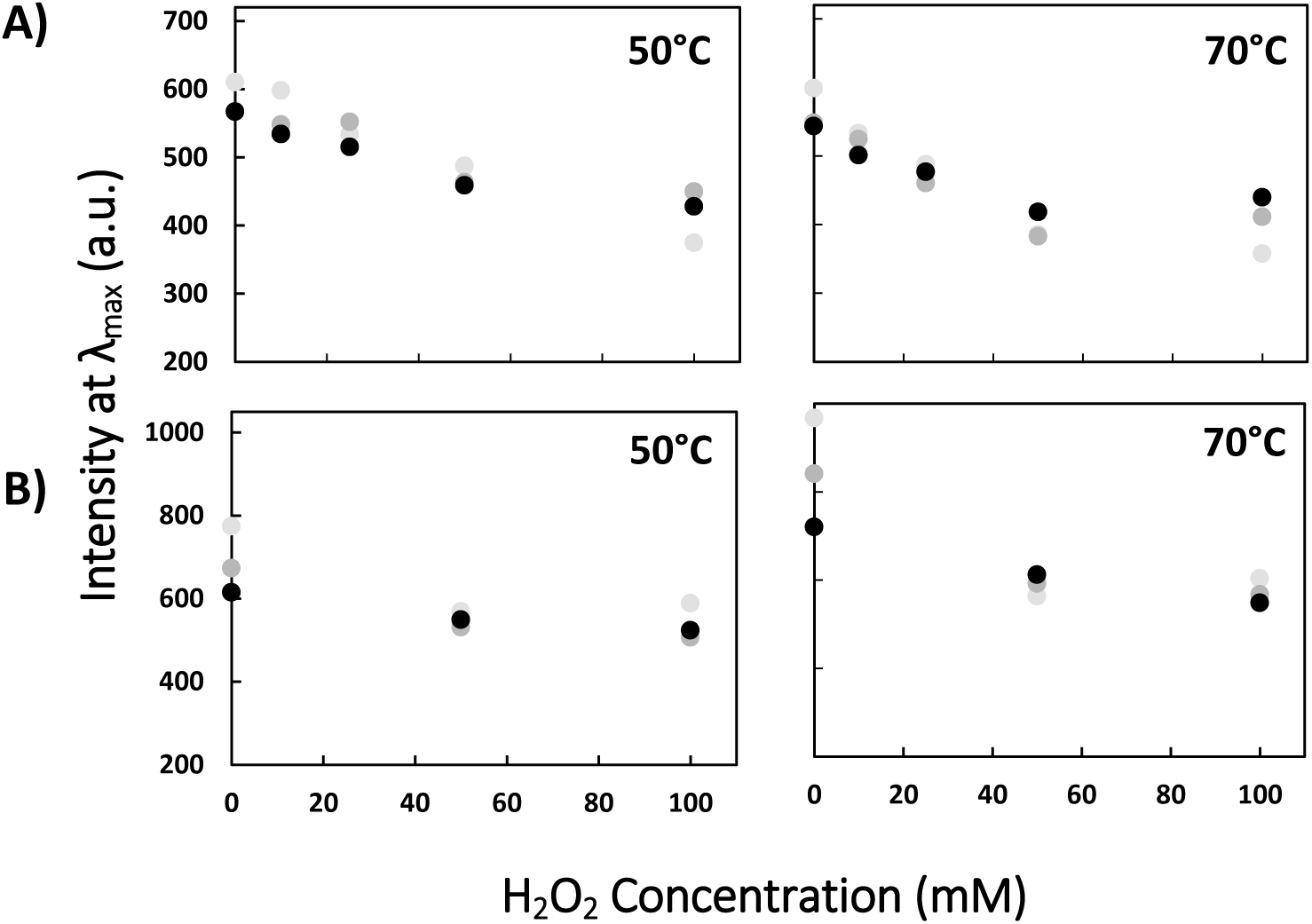
Structural changes of artemin under different stress conditions are irreversible. A) Artemin at final concentration of 0.1 mg/mL was treated with 0-100 mM H_2_O_2_ for 6 h at 0°C in dark, followed by incubating at 50 and 70°C for 20 min. Then the samples were kept at 12°C for 48 hours and the Trp fluorescence intensity was recorded with the excitation wavelength at 280 nm and the fluorescence intensity at λ_max_ was determined. B) ANS (30 µM) fluorescence intensity at λ_max_ in the presence of 0.25 mg/mL HO-artemin with the excitation wavelength at 380 nm.

### Chaperone-like activity of artemin under heat and oxidative stress

#### H-artemin

Denatured*/*reduced lysozyme was refolded by the dilution method and the kinetics of chaperone-assisted refolding was examined in the presence of 0.5 and 1 µg/mL artemin (Figure 9). As shown in Figure 9A, B, the heated artemin at 25 and 50°C was found efficient in suppressing aggregation of the enzyme. In contrast, the 0.5 µg/mL chaperone incubated at elevated temperatures, 60 to 80°C, accelerated the aggregation of lysozyme (Figure 9A), and 1 µg/mL artemin showed no effect on the refolding yield at similar conditions (Figure 9B).

**Figure 9.**
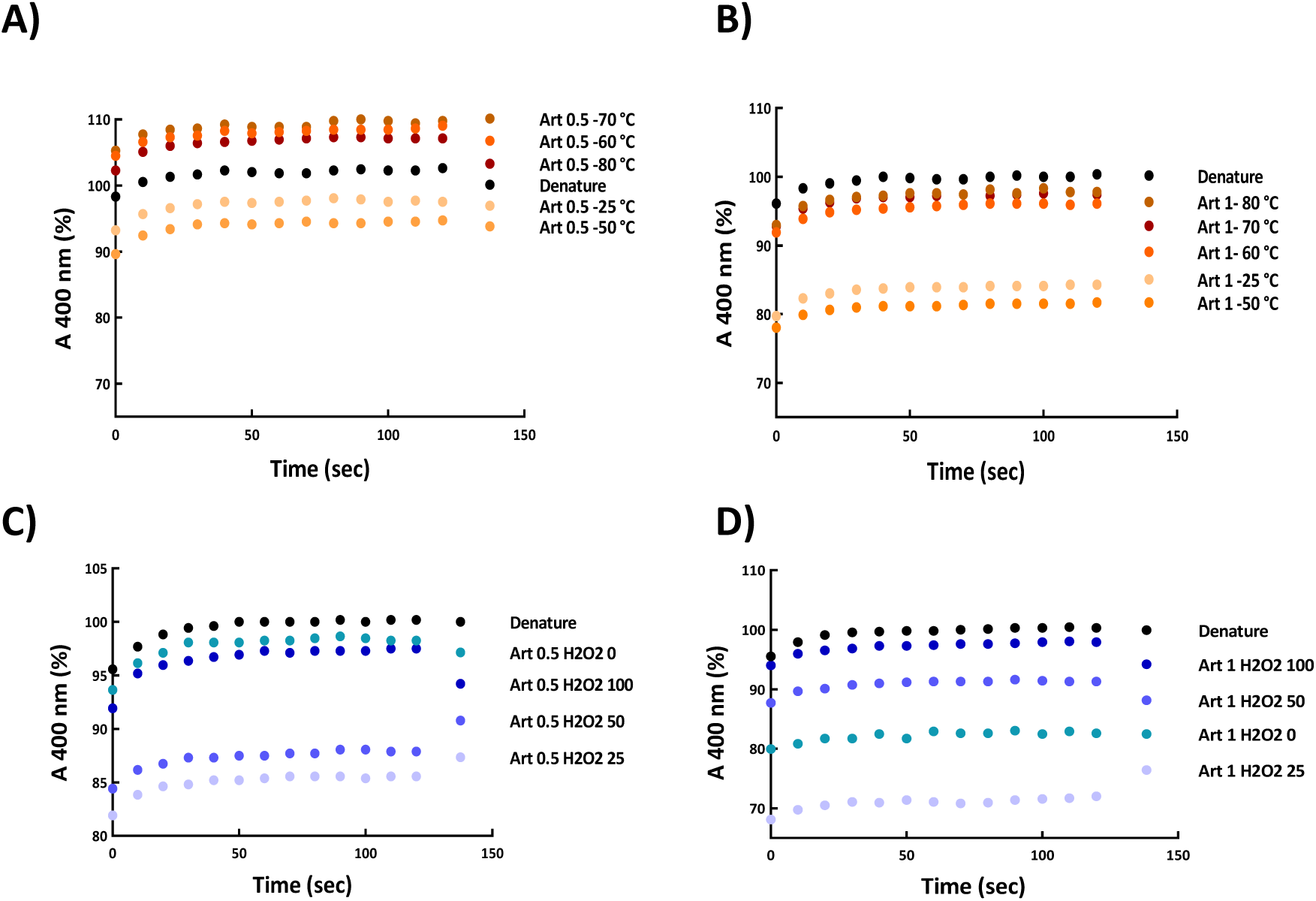
Refolding of lysozyme in the absence (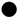) and presence of (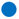) heated (A, B) and (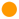) oxidized (C, D) artemin. Denatured lysozyme (10 mg/mL) in a solution containing 40 mM DTT, 6 M GdmCl, and 50 mM potassium phosphate buffer, pH 7.1, was diluted with a mixing ratio of 1:50 by the refolding buffer. The diluted sample contained 0.2 mg/mL lysozyme, 0.8 mM DTT, 0.12 mM GdmCl and 50 mM potassium phosphate buffer, pH 8.5, 5 mM GSH, 5 mM GSSG, and 0.5 (A, C) and 1 µg/mL (B, D) artemin. The kinetic of refolding was recorded by monitoring light scattering at 400 nm at 25°C.

#### O-artemin

Results showed that the oxidized chaperone (0.5 µg/mL) with 25 and 50 mM H_2_O_2_ was efficient in preventing the aggregation of lysozyme compared to the untreated control (Figure 9C). In contrast, the anti-aggregatory potency of O-artemin decreased by increasing the oxidant content, and the oxidized chaperone with 100 mM H_2_O_2_ did not show any effect on the enzyme refolding yield (Figure 9B, C).

#### HO-artemin

Figure 10 depicts the impact of HO-artemin (1 µg/mL) on refolding of lysozyme based on constant concentrations of H_2_O_2_ (25, 50 and 100 mM) (Figure 10A, B, C) and temperatures (25, 50 and 70°C) (Figure 10D, E, F). According to Figure 10A, the HO-artemin at 25 and 50°C suppressed the aggregation of lysozyme, but at elevated temperature, 70°C, it could not efficiently prevent the aggregation of lysozyme (Figure 10A, B). These trends were similar to the graphs obtained for H-artemin (Figure 9 A,B). In contrast, HO-artemin incubated with 100 mM H_2_O_2_ clearly exhibited an improved chaperone activity (Figure 10C).

**Figure 10.**
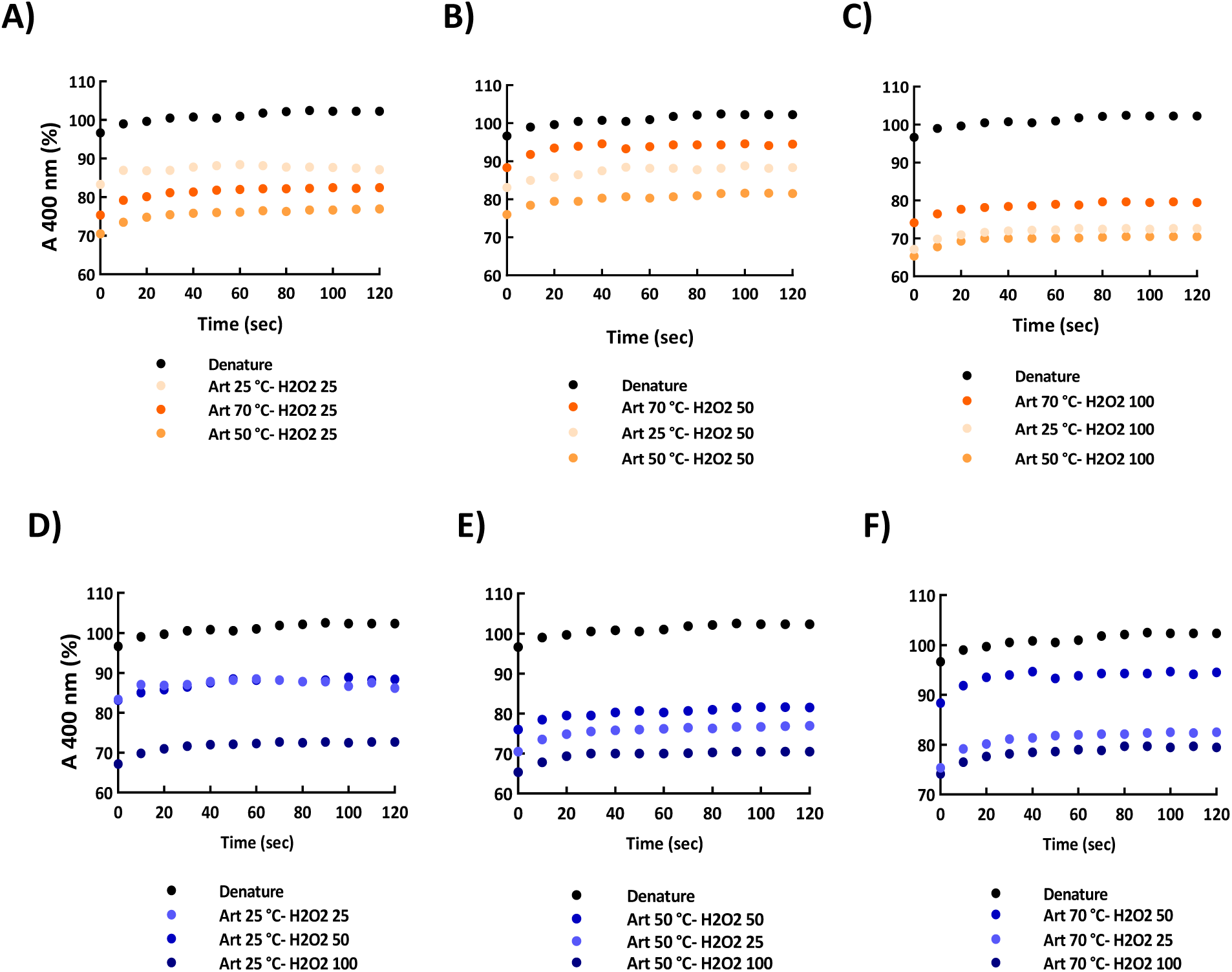
Influence of heated-oxidized artemin on aggregation accompanying refolding of lysozyme. The comparison was performed based on constant (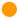) H_2_O_2_ concentrations (A-C) and (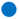) temperatures (D-F). Artemin at final concentration of 1 µg/mL was used and the denatured sample without added chaperone was introduced as control (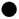). The kinetic of refolding was recorded by monitoring light scattering at 400 nm at 25°C.

We also checked the effect of HO-artemin on refolding of lysozyme based on constant temperatures (Figure 10D, E, F). Results clearly demonstrated an enhanced capacity of the chaperone along with the increased oxidant content at 25°C (Figure 10A). At higher temperatures (50 and 70°C), oxidized artemin with 50 and 100 mM H_2_O_2_ showed a similar potency (Figure 10B, C. Comparing these observations with the results from the effect of O-artemin on refolding yield (Figure 9C, D) reveals that the chaperoning function of HO-artemin was considerably improved in presence of both stressors.

## DISCUSSION

Artemin is an abundant heat stable protein in *Artemia* encysted embryos and it was found that high regulatory production of artemin under harsh environmental conditions is probably relevant to stress resistance in this crustacean (2, 16). Artemin demonstrated an ability in suppressing heat-induced aggregation of different protein substrates such as citrate synthase, carbonic anhydrase, horseradish peroxidase and luciferase *in vitro* and *in vivo*, and also introduced as a potent nucleic acid chaperone (6–9,11). The intrinsic conformational properties of artemin seem to play a critical role in its biological activities. Our previous structural report documented that artemin contains a high surface hydrophobicity compared to other molecular chaperones (9). Moreover, the chaperone potency of artemin is greatly dependent on the presence of cysteine residues and intermolecular disulfide bond formation (13). Accordingly, heat and oxidation are among the most important factors which probably affect the chaperone capacity of artemin. In this report, we have evaluated the conformational changes of artemin under heat and oxidative stress using structural and functional analysis. The results have been summarized in Table 1. Our findings may help us in better understanding various mechanisms of sHsps in protecting clients in different stresses.

**Table 1.** A summarize of the structural and functional changes of artemin upon heat and H_2_O_2_ treatments.

| **Stress**<br>**Changes** | **Heat** | **H<sub>2</sub>O<sub>2</sub>** | **Heat & H<sub>2</sub>O<sub>2</sub>** |
| --- | --- | --- | --- |
| **Hydrophobicity** | 70-80°C: ↑ | 0-40 mM: ↓<br>80-160 mM: ↑ | 50, 70°C; 10-100 mM: ↓ |
| **Dimerization** | 25-50°C: ↑<br>60-80°C: ↓ | 0-40 mM: ↑<br>80-160 mM: ↓ | 50, 70°C; 10-100 mM: ↑ |
| **Reversibility** | Irreversible | Irreversible | Irreversible |
| **Chaperone Activity** | 25-50°C: Less to More<br>50-80°C: More to Less | 0-25 mM: Less to More<br>50-100 mM: More to Less | 50-70°C; 25-100 mM:<br>Less to More |

There are different mechanisms for small heat shock proteins (sHsps) to protect protein substrates in different stresses (17). Many sHsps are known to undergo temperature-dependent changes in their structures, which modulate their chaperone activity (18). These mechanisms may be mediated through conformational transitions between dimerization and oligomerization behavior of the chaperones. For example, at elevated temperatures, dissociation of large oligomeric forms of Hsp26 into smaller, active species of the chaperone occurs (19) and also temperature-dependent rearrangements in the tertiary structure have been reported for Hsp22 which improve its chaperone activity (20). Hsp33 is a redox-regulated chaperone whose activity is mainly mediated by the heat effect and inducing its dimerization (21). Besides, Hsp20.1 and Hsp14.1 oligomers dissociated to smaller oligomeric forms or even dimer/monomer species under acid stresses (17). These dissociations seem to be necessary for the exposure of additional hydrophobic sites on the surface of the protein molecule (22).

### Heat-dependent structural transitions

It is expected that at elevated temperatures, artemin undergoes heat-dependent structural transitions as well as other sHsps to expose a large amount of hydrophobic residues on surfaces of the protein. Because artemin contains seven Trp and five Tyr residues, monitoring its conformational alteration was possible using its intrinsic fluorescence. At present, there is no reliable evidence for the accurate spatial localization of these amino acids in artemin. It has been confirmed that Trp and Tyr fluorescence is usually quenched in a distance-dependent manner. Accordingly, when such aromatic amino acids are located on the surface of the protein, the fluorescence intensity of the residues is usually influenced in a higher degree by the quencher, compared to the amino acids deeply embedded in the protein structure (23). Our intrinsic fluorescence results showed that the fluorescence emission of the protein was significantly reduced at elevated temperatures and the first transition observed at 50°C followed by the second transition at 80°C (Figure 1A). This is suggested that the induced conformational changes of artemin resulted in the increased quenching probably due to the exposing internal aromatic residues to solvent.

The conformational changes of the protein was also investigated using ANS fluorescence. ANS is widely used as a fluorescent hydrophobic probe for hydrophobic patches of proteins (22, 24). It is basically non-fluorescent in aqueous solutions but became highly fluorescent in non-polar environment (22). Increases in temperature up to 60°C caused artemin to present more exposed non-polar sites and binding ANS to its hydrophobic regions resulting in an enhanced fluorescence intensity and a weak blue shift in the λ_max_ (Figure 1B) as also illustrated by other reports (22, 25). Such hydrophobic interactions in artemin likely play a critical role in defining conformation and mediating the protein-chaperone interactions (24). In contrast, the fluorescence intensity of the incubated protein at lower temperatures did not change which presumably reflects low changes in surface hydrophobicity of the heated proteins, particularly the lack of ANS-binding sites. As a point, due to the thermal inactivation of the excited state of the probe, the heat-treated proteins allowed to cool down at room temperature for 20 min followed by fluorescence measurements (24).

In a second approach, different monomer, dimer and oligomer forms of H-artemin were detected under thermal conditions using SDS-PAGE technique after addition of 0.5% glutaraldehyde as a fixative agent. Glutaraldehyde cross-linking is a method normally used for determination of the subunit structure of oligomeric proteins (26). SDS-PAGE result showed that the uncross-linked protein was mainly presented as monomers at 25°C (Figure 2A), but the dimer population found to be dominated under cross-linking condition. The abundance of the 54 kDa protein band was slightly enhanced upon increasing temperatures from 25 to 50°C, but became significantly weaker at 60°C and completely disappeared at 70 and 80°C, probably due to the formation of higher molecular weight oligomers and/ or aggregates by hydrophobic interactions that could not enter the gel (Figure 2A). In addition, we examined whether the heat-induced dimerization mediates *via* disulfide bonds by addition of DTT as a reducing agent to the protein solutions at the concentrations used for reducing the chaperone before the heat treatments. Accordingly, the dimers were disappeared in the presence of DTT confirming that the disulfide bond(s) are necessary for dimerization (Figure 2B). The size distribution analysis of H-artemin showed that the size of proteins was enhanced upon increasing temperatures probably due to formation of dimers, oligomers (Figure 3A,B) and probably protein oligomers and/ or aggregates (Figure 3C, D) as a result of hydrophobic interactions (Figure 1B).

According to many previous reports, sHsps generally use their N-terminus to bind client proteins due to the presence of a lot of hydrophobic residues in this region which is located on the inner surface of the oligomers (27). So, heat-induced oligomer dissociation may expose a large number of hydrophobic surfaces that are normally hidden inside the oligomer, including not only the N-terminus, but also the dimer–dimer and even the monomer–monomer interfaces (17). As a result, the substrate-binding ability of the chaperones would be improved in a temperature-dependent manner to cope with the increased client substrates within cells (28). Totally, our results revealed that at elevated temperatures, H-artemin presumably undergoes the structural changes which are associated with a marked increase in the surface hydrophobicity and also degree of dimerization of the chaperone *via* disulfide bridges in order to improve the binding capacity of the chaperone to the client proteins in an unfolding intermediate state. As a suggestion, interaction of the protein substrates may subsequently prevent the molecular chaperone with a highly exposed hydrophobic surface from self-association and participation during the stress conditions.

### Oxidative-dependent structural transitions

Artemin contains a high content of cysteine residues and due to the high cysteine numbers and their specific distribution, they may be regarded as critical factors in regulating chaperone function (1, 13). We have shown earlier that oxidation/reduction of cysteine residues influences chaperone potency of artemin (13). Experimental results revealed that 9 out of 10 thiols are free in artemin monomers and there is only one cysteine involved in inter-molecular disulfide bond formation. Molecular modeling also predicted Cys22 with the highest accessible surface area, which is probably the responsible residue for inter-subunit disulfide bond formation (13).

Accordingly, it was supposed that two monomers of artemin form the dimer through an inter-molecular disulfide bond and results in the protein switches between its less and more active forms. Here we tried to provide detailed structural information on conformational changes of artemin during oxidative conditions. Intrinsic fluorescence measurements demonstrated that upon increasing hydrogen peroxide concentration to 40 mM, artemin structure was still stable at these concentrations of the oxidant (Figure 4A, Figure 5), despite many other susceptible proteins (29, 30). However, higher concentrations of the oxidant (80-160 mM) led to a remarkable reduction in the fluorescence intensity of O-artemin with a slight shift of λ_max_ to longer wavelengths (Figure 4A, insert) which showed that oxidation changed the microenvironment around the aromatic residues. It is well known that Trp and Tyr residues have a preference for negative charge residues, cysteine and disulfide bonds in their spatial environments, and the interactions of sulfur atoms with the aromatic rings may lead to the fluorescence decay (31). Cystines in disulfide bonds have a high propensity to interact with tryptophan residues (32). The breakage or formation of the disulfide bonds can induce fluorescence changes of proteins as a result of modifications of the micro-environment of the fluorescent residues (33). Accordingly, variable degrees of fluorescence quenching of Trp and Tyr residues probably occurred by nearby formed disulfide and sulfhydryl groups in O-artemin (Figure 4A).

ANS binding indicated that increasing H_2_O_2_ content from 10 to 40 mM led to a decline in fluorescence intensity (Figure 4B), as well as a shift in λ_max_ to longer wavelengths (Figure 4B, insert). Therefore, it seems that the surface hydrophobicity of O-artemin has been relatively decreased. It is suggested that disulfide bond formation due to oxidation of Cys residues resulted in oligomerization of artemin and at the same time hydrophobic patches were buried deeply within the protein oligomer assemblies. Although, higher concentrations of the oxidant (80-160 mM) resulted in an increase in ANS binding, the increase was not significant as well as for H-artemin (Figure 1B). Similar trends were also recorded for the chaperone GroEL in the presence of H_2_O_2_ (34).

In order to further confirm the dimerization/oligomerization status of O-artemin, we performed glutaraldehyde cross-linking of artemin oxidized in the presence of 0-160 mM H_2_O_2_. SDS-PAGE illustrated that along with increasing the oxidant content from 0 to 100 mM H_2_O_2_, the 54 kD dimeric bands gradually strengthened (Figure 5) and the bands weakened at 160 mM H_2_O_2_, probably due to the formation of a higher degree of oligomers. It is hypothesized that the highly reactive thiol groups of cysteine residues may lead to protein oligomerization due to the excess formation of disulfide bonds in O-artemin. Finally, the protein precipitation may occur as a result of highly exposed hydrophobic regions and the formation of excess inter-protein disulfide bridges (32) as showed by fluorescence (Figure 4B) and SDS-PAGE analysis (Figure 5). Totally, these results indicate that upon oxidation, artemin undergoes dimerization and oligomerization.

### Structural transitions of artemin under heat and oxidant treatments

In a final approach, we also examined the simultaneous effect of stressors on protein structure. Artemin was exposed to different concentrations of H_2_O_2_ (0-100 mM), followed by incubation at elevated temperatures (50, 70°C). The fluorescence measurements showed that intensity decreased upon increasing the oxidant content at both temperatures (Figure 6A). This result is in parallel with fluorescence measurement of O-artemin (Figure 4A). ANS binding also showed a decreasing trend of the fluorescence intensity along with the increased oxidant concentrations up to 100 mM H_2_O_2_ for HO-artemin (Figure 6B), although O-artemin showed an increased ANS binding upon treatment with 80-160 mM H_2_O_2_ (Figure 4B). Our suggestion is that the oxidant agent sequestered the exposed hydrophobic surfaces on the heated chaperone by stabilizing the dimeric forms and preventing the protein from aggregation. Besides, we checked the tertiary structural changes of HO-artemin using protein cross-linking analysis by SDS-PAGE (Figure 7). In agreement with the results obtained for O-artemin (Figure 5) and H-artemin (Figure 2A), the dimerization of HO-artemin was enhanced upon increasing the oxidant concentrations from 0-100 mM H_2_O_2_ at both temperature treatments and the degree of dimerization was higher at 50°C in comparison with 70°C (Figure 7).

The reversibility of the structural changes of artemin was investigated using fluorescence measurements (Figure 8). Both intrinsic (Figure 8A) and extrinsic (Figure 8B) fluorescence analysis showed that artemin structure altered irreversibly under heat and oxidative conditions.

### Heat/Oxidative-dependent chaperone activity

To assess the potency of heated/oxidized artemin on the aggregation accompanying refolding of lysozyme, refolding experiments were done in the presence of 0.5 and 1 µg/mL artemin. Our results indicated that H-artemin incubated at 25 and 50°C promoted the refolding of the enzyme (Figure 9A, B), however, at elevated temperatures (60-80°C), it accelerated the enzyme aggregation (Figure 9A). In addition, oxidation with lower concentrations of H_2_O_2_ (25, 50 mM) activated the chaperone and subsequently promoted the refolding of the enzyme, but, the higher degree of oxidation of artemin (100 mM H_2_O_2_) did not affect the lysozyme refolding yield (Figure 9C, D). Our suggestion is that higher concentrations of the oxidant influenced the structure of artemin and a weak chaperone activity observed as a result of oxidative modification of some amino acids of the protein, and changing the quality of hydrophobic groups of the protein (34) which may lead to self-association/agglomeration of the chaperone as also confirmed by fluorescence (Figure 4) and protein cross-link (Figure 5) analysis.

Finally, we performed chaperone activity assays using HO-artemin. The results exhibited that the ability of HO-artemin in preventing the enzyme aggregation was significantly improved (Figure 10). Analysis of data revealed that the oxidant did not strongly influence the heated chaperone (Figure 10A,B). The results suggested that there is a similar anti-aggregatory trend for H-artemin and HO-artemin, especially at 25 and 50 mM H_2_O_2_ (Figure 9B and Figure 10A-C). In contrast, the HO-artemin exhibited a higher efficacy in suppressing the enzyme aggregation (Figure 10D-F) compared to O-artemin (50 and 100 mM H_2_O_2_) (Figure 9D). Our observations revealed that although the presence of both heat and oxidant is essential to achieve the most active form of the chaperone, heat has a stronger impact on the structure and function of artemin than the oxidizing agent.

Here, we reported that the mechanism of artemin’s activation mainly relies on dimerization of the protein monomers which is strongly modulated through heat-induced conformational changes of the protein. Besides, such dimer and/or oligomer structures can be stabilized through the formation of inter-subunit disulfide bridge(s) (12) in the dimer form and switch the less active to the more active form of proteins. Subsequently, stable dimerization of the artemin monomers leads to accumulation of highly active artemin species. It has been indicated that disulfide bonds improve the thermal stability of proteins with an activity in the oxidizing extracellular environment (32). One explanation is that disulfide bonds reduce the conformational freedom and entropy of the protein in its unfolded state, and consequently destabilize this state with respect to the folded state (35). Similar mechanisms have been proposed for the other chaperone, Hsp33 and Hsp27 which control their dimerization by forming an intermolecular disulfide bond (21, 36). Our observations suggested that there may be different mechanisms for artemin to protect client proteins in different stresses which are basically mediated through protein’s dimerization (Scheme 1). The proposed mechanisms presumably play a vital role in conserving the *Artemia* cysts’s tolerance against severe environmental stresses.

**Scheme 1.**
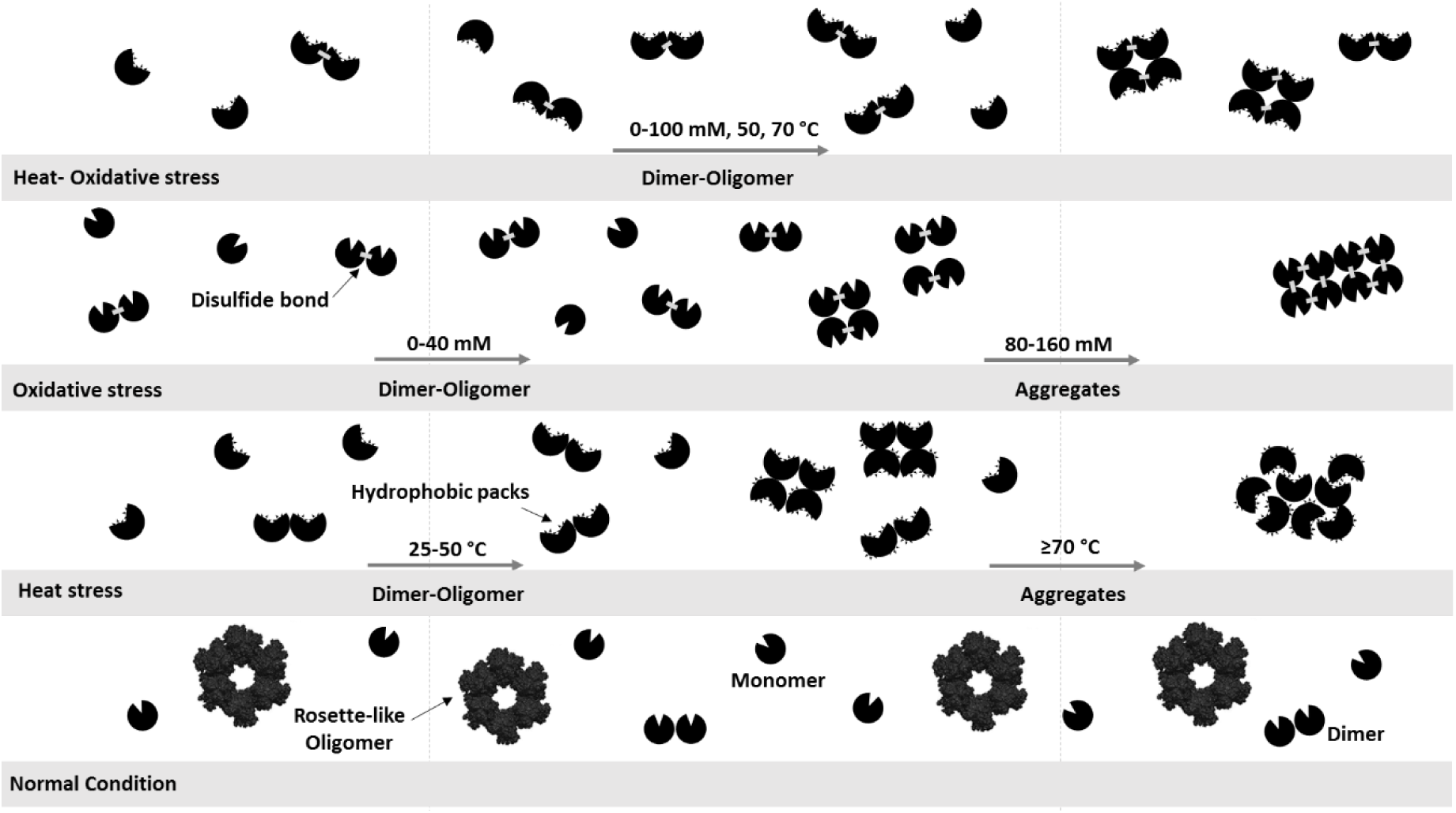
Model of chaperone function of artemin. Under physiological condition, artemin exists mainly in rosette-like oligomeric forms. Upon heat shock, **t**he exposed hydrophobic packets of dimeric structures of the chaperone probably play a vital role in mediating the protein-chaperone interactions. At elevated temperatures, the highly exposed hydrophobic surfaces of the chaperone lead to its self-assembly as aggregates. The oxidized artemin also forms stable dimers through formation of disulfide bridges between the chaperone monomers and the protein oligomerization/agglomeration occurs as a consequent of the exposure of a high degree of hydrophobic surfaces and the formation of inter-protein disulfide bridges. Under both stress conditions, the most active stable form of the chaperone is achieved through formation of the stable dimers with an appropriate exposed hydrophobic sites. Through these mechanisms, artemin oligomers dissociate into dimers and monomers upon heat and/or oxidative stresses, which are able to bind non-native proteins, thus preventing their aggregation.

Our finding confirmed that artemin appears to exist in different conformational forms including monomer, dimer and oligomer as a function of heat and oxidant. Both hydrophobicity and dimerization are important to achieve the chaperone in a fully active form. At elevated temperatures and higher degree of oxidation, both hydrophobicity and dimerization increased. In contrast, under both stress conditions, hydrophobicity did not change at the higher oxidant concentrations, while the dimerization was enhanced. Such dimerization strongly relied on disulfide bond formations between the artemin monomers. The protein cross-link experiments indicated that the oligomerization of artemin as a consequence of self-association at elevated temperatures. Our suggestion is that upon increasing temperature up to 60°C, artemin exposes the maximum hydrophobic sites in order to stably bind the target folding intermediates. Therefore, the presence of such binding substrates may stabilize artemin conformation at higher temperatures and protect it from self-association/precipitation and this may result in the reversible structural changes of artemin upon exposing to stress conditions. Enhanced peptide*-* substrate binding upon heat treatment was previously reported for other molecular chaperones Hsp26 and gp96 (19, 37). In the case of gp96, it was recognized that the heat-induced oligomers retain peptide binding ability and it was suggested that these soluble aggregates could serve as a reservoir, and be converted into activated chaperone molecules under certain circumstances (38). Our aggregation accompanying refolding experiments also showed the chaperone potency was considerably influenced by the temperature rather than the oxidant. Also, the artemin function was improved at the lower oxidant concentrations probably due to proper dimerization of the chaperone through disulfide bond formation as it was detected by cross-link analysis. Despite the weak chaperone potency of oxidized artemin with high concentrations of H_2_O_2_, simultaneous incubation of the chaperone with elevated temperatures and oxidant concentrations significantly triggered the activation of chaperone. As the fluorescence studies showed, the presence of the oxidant resulted in a lower ANS binding through sequestering the exposed hydrophobic patches on the heated artemin probably to inhibit its aggregation. It is suggested that oxidation presumably acts through stabilizing the dimer structures of artemin through formation of disulfide bridges between the protein monomers and strengthens its stability and chaperoning potency.

## CONCLUSION

It can be concluded that the heat-induced dimerization of artemin is the most critical factor for its activation. The heat and oxidation are important stresses which *Artemia* cysts face in their environments. Totally, it seems that *in vivo* cytosolic artemin may exist in a monomer–oligomer equilibrium in the cytoplasm of the cysts which play a critical role in maintaining the cysts under such extreme conditions. Environmental stresses and/or intracellular portion of proteins may shift the equilibrium towards the active dimer forms. Although the function of artemin has been thoroughly investigated *in vitro* and *in vivo,* its mechanism is not yet completely examined. This work has presented some evidence in support of the relation between artemin and its heat/oxidation-dependent activation *in vitro*. Further studies can be also performed in future for illustrating the substrate-chaperone interactions in stress conditions.

## AUTHOR CONTRIBUTIONS

ZT performed the experiments, analyzed and interpreted data and contributed to manuscript writing. ZA was involved in performing the experiments. BM contributed to the spectroscopic methods. SSS was involved in the formation of the experimental concept and revised the manuscript. RHS proposed the original idea, designed and supervised the experiments, provided intellectual input, and edited the manuscript. ‡These authors contributed equally.

## ACKNOWLEDGMENTS

This work was supported by the research council of Tarbiat Modares University and Ministry of Sciences, Researches, and Technology, Iran.

